# New strains for tissue-specific RNAi studies in *Caenorhabditis elegans*

**DOI:** 10.1101/283325

**Authors:** Jason S. Watts, Henry F. Harrison, Shizue Omi, Quentin Guenthers, James Dalelio, Nathalie Pujol, Jennifer L. Watts

**Author notes:** Author for correspondence: Jennifer L. Watts, School of Molecular Biosciences, Washington State University, PO Box 647520, Pullman, WA 99164-7520.

## Abstract

RNA interference is a powerful tool for dissecting gene function. In *Caenorhabditis elegans*, ingestion of double stranded RNA causes strong, systemic knockdown of target genes. Further insight into gene function can be revealed by tissue-specific RNAi techniques. Currently available tissue-specific *C. elegans* strains rely on rescue of RNAi function in a desired tissue or cell in an otherwise RNAi deficient genetic background. We attempted to assess the contribution of specific tissues to polyunsaturated fatty acid (PUFA) synthesis using currently available tissue-specific RNAi strains. We discovered that *rde-1 (ne219)*, a commonly used RNAi-resistant mutant strain, retains considerable RNAi capacity against RNAi directed at PUFA synthesis genes. By measuring changes in the fatty acid products of the desaturase enzymes that synthesize PUFAs, we found that the before mentioned strain, *rde-1 (ne219)* and the reported germline only RNAi strain, *rrf-1 (pk1417)* are not appropriate genetic backgrounds for tissue-specific RNAi experiments. However, the knockout mutant *rde-1 (ne300)* was strongly resistant to dsRNA induced RNAi, and thus is more appropriate for construction of a robust tissue-specific RNAi strains. Using newly constructed strains in the *rde-1(null)* background, we found considerable desaturase activity in intestinal, epidermal, and germline tissues, but not in muscle. The RNAi-specific strains reported in this study will be useful tools for *C. elegans* researchers studying a variety of biological processes.

## Introduction

RNA interference (RNAi) is an evolutionarily ancient defense mechanism against viruses and transposable elements. *Caenorhabditis elegans* has been a powerful system for discovering the molecular mechanisms underlying the RNAi phenomena and its role in gene regulation (Fire et al. 1998; Grishok 2005; Hammond 2005). The discovery that double stranded RNA (dsRNA) expressed by bacteria and ingested by the worm could effectively silence target genes revolutionized the use of RNAi as a tool for high-throughput, large-scale genetic knockdown studies in *C. elegans* (Timmons and Fire 1998; Fraser et al. 2000; Ashrafi et al. 2003; Yigit et al. 2006).

Genetic screens for mutants that are resistant to RNAi have been essential for elucidating the mechanism of processing exogenously introduced double stranded RNAs and inducing gene silencing (Zhuang and Hunter 2012). The feeding method of RNAi delivery is successful in *C. elegans* because of two membrane proteins called SID-1 and SID-2 that facilitate the uptake of double stranded RNA into cells (Winston et al. 2002; Winston et al. 2007). Both mutant strains grow and reproduce normally but are resistant to systematic RNAi delivered by the feeding method. Other screens for viable mutants with RNAi deficiency (RDE mutants) revealed genes coding for many highly-conserved activities required for RNAi, including RDE-4, a double-stranded RNA binding protein which forms a complex with Dicer to bind long double-stranded RNAi and cleave it into ~22 bp interfering RNAs (siRNAs) (Tabara et al. 2002; Knight and Bass 2001; Parker et al. 2006).

In *C. elegans*, RDE-1 is the primary Argonaute component of the RNA induced silencing complex (RISC), which degrades the passenger strand of the siRNA and uses the remaining strand to target mRNA for use as a template for synthesis of secondary siRNAs (Tabara et al. 1999; Parrish and Fire 2001; Steiner et al. 2009). Production of the siRNAs amplifies the signal and are used as the final targeting signal for degradation of newly formed mRNAs. The *C. elegans* genome encodes multiple homologs of certain components of the RNAi machinery, including four RNA-dependent RNA polymerases (RdRPs), including RRF-1, and 27 known Argonaute proteins (Smardon et al. 2000; Sijen et al. 2001; Yigit et al. 2006). The RdRPs amplify silencing by using the primary siRNA/mRNA complex as a template for synthesizing secondary siRNAs. RDE-3, a member of the polymerase beta nucleotidyltransferase superfamily is required for siRNA accumulation (Chen et al. 2005). RDE-10 and RDE-11 form a complex that promotes secondary siRNA amplification (Yang et al. 2012).

Identifying mutants that are resistant to the induction of RNAi by double stranded RNA has typically involved screening with a visible phenotype such as lethality, movement defects, or suppression of a fluorescent transgene (Tabara et al. 1999; Yigit et al. 2006; Winston et al. 2002). Mutant strains in which the treated worms failed to display the knockdown phenotype were considered RNAi resistant. These methods have proven efficient and invaluable for the discovery of RNAi pathway genes. However, the qualitative nature of these visible phenotypes makes them insufficient to determine if a specific mutation in an RNAi pathway gene completely inhibits RNAi capacity, or only enough to prevent the visible phenotype.

Gene knockdown by RNAi is useful for elucidating gene function, but it does not provide information about the function of genes in specific tissues. To assess tissue specific gene function, investigators have employed tissue specific transgenic rescue of key RNAi machinery (Sijen et al. 2001; Qadota et al. 2007). Accurate analysis of experiments performed in tissue specific systems depends on RNAi functioning only in the desired cells and tissues. This is achieved by expressing a transgene under the control of a tissue-specific promoter in the background of an RNAi-deficient mutant. Any residual RNAi activity in off-target tissues can lead to erroneous conclusions, thus it is essential that the mutant strain used is completely RNAi-deficient.

Our objective was to determine various tissue contributions to the overall degree of fatty acid desaturation in the nematode *C. elegans* using tissue-specific RNAi strains and feeding RNAi directed toward two fatty acid desaturase genes. We used a biochemical method employing gas chromatography/mass spectrometry (GC/MS) to monitor the flux of fatty acid desaturation reactions from substrates to products as a quantitative approach to measuring the RNAi silencing capacity in previously characterized *C. elegans* mutant strains. Our quantitative method revealed that some *C. elegans* strains deemed “RNAi deficient” retained substantial RNAi silencing capacity when exposed to the feeding method of RNAi delivery, and therefore the available strains used for tissue-specific RNAi are not useful for determining tissue-specificity of gene function. We present here a collection of new strains for tissue-specific RNAi and report that intestine and epidermal tissues produce the highest amounts of desaturated fatty acids, while the germline also shows activity of omega-3 and delta-5 fatty acid desaturases.

## Results

### Using polyunsaturated fatty acid desaturation to assess RNAi efficiency

Fatty acids are the building blocks of the lipids that cells use for energy storage, membrane structure, and signaling. *C. elegans* obtains fatty acids from its bacterial diet, and also has the capacity to generate fatty acids *de novo* from acetyl-CoA (Watts and Ristow 2017). Saturated fatty acids obtained from the diet or synthesized *de novo* are further modified by chain elongation and the addition of one or more double bonds, (Watts and Browse 2002) greatly enhancing the structural diversity and utility of fatty acids available to the cell. Based on published RNA sequencing studies (Chikina et al. 2009; Cao et al. 2017) and on reporter strains constructed in our lab, we suspected that fatty acid desaturases are expressed in various tissues in *C. elegans*, with the highest expression in the intestine, the tissue responsible for absorption of dietary fats and modification of synthesized and dietary fats for incorporation into membrane lipids, and as well as for storage lipids contained in lipid droplets and yolk (Watts and Ristow 2017). We reasoned that using tissue specific RNAi strains, and determination of fatty acid composition by GC/MS, we could establish the relative contributions of various tissues to overall fatty acid composition.

We focused on two of the FA desaturase enzymes, FAT-1, an omega-3 desaturase, and FAT-4, a Δ5 desaturase (Watts and Browse 2002). FAT-1 adds double bonds between the third and fourth carbons of a FA from the methyl end (omega-3), whereas the FAT-4 Δ5 desaturase adds double bonds between the fifth and sixth carbon counting from the carboxyl end of the fatty acid. **Figure 1** shows the fatty acid desaturation pathway, highlighting the substrates used by, and products synthesized by FAT-1 (**Figure 1A**) and FAT-4 desaturases (**Figure 1C**). *C. elegans* grown on bacteria expressing double stranded RNA complementary to *fat-1* or *fat-4* grow normally and reproduce, as do *fat-1* and *fat-4* null mutants (Watts and Browse 2002).

**Figure 1.**
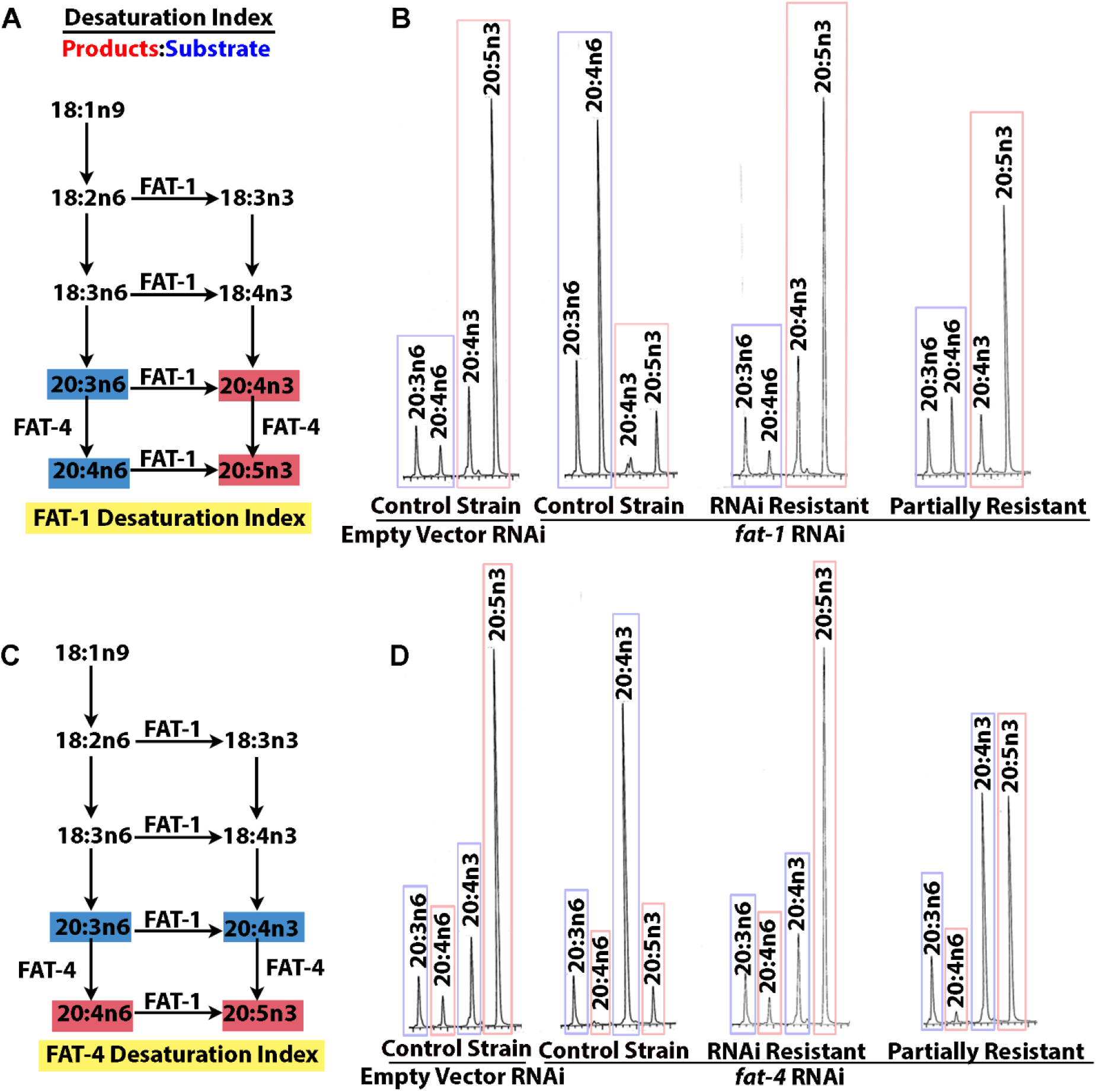
Determination of desaturation index. To measure RNAi deficiency in mutant worm strains, we knocked down the fatty acid desaturase genes *fat-1* and *fat-4* with RNAi by feeding and determined the ratio of fatty acid desaturation products (red) to their substrates (blue) and termed this the desaturation index for each desaturation reaction. The example chromatographs demonstrate that in RNAi competent worms (control strain), treatment with RNAi against either *fat-1* (A) or *fat-4* (D) causes accumulation of substrates (blue) and depletion of products (red). Example chromatographs of and RNAi “resistant strain” show a high ratio of products to substrates (Desaturation index), similar to empty vector control treated worms, and chromatographs of the “partially resistant” strain have an intermediate desaturation index.

Loss of function mutations and RNAi treatment of *fat-1* and *fat-4* lead to reduced ratios of specific fatty acid products to substrates, a relationship that can be quantified as the desaturation index. When grown in lab conditions, on plates with *E. coli* HT115 bacteria, wild type *C. elegans* typically shows a desaturation index of 4-6 corresponding to FAT-1 activity, meaning the abundance of omega-3 fatty acid products is 4-6 fold more than the omega-6 substrates. For FAT-4, the desaturation index is typically 2.5-3, meaning the Δ5 desaturated products are 2.5-3 fold more abundant than their substrates (see Materials and Methods for desaturase index calculations). In *fat-1* or *fat-4* null mutants, the corresponding desaturation indices drop to zero because there are no omega-3 or delta-5 unsaturated PUFAs formed in the mutant strains. Feeding RNAi is a very efficient means to knock down FAT-1 and FAT-4 activity, and in a wild type background, *fat-1* RNAi causes a drop in the desaturation index from 4-6 to 0.2. Similarly, *fat-4* RNAi causes a drop in the desaturation index of 2.5-3 down to 0.1 (**Figure 1 and Table 1**). An RNAi-resistant strain treated with *fat-1* or *fat-4* RNAi would be expected to show a similar desaturation index as an untreated strain. **Figure 1B and 1D** show typical chromatographs of the wild type control strain, as well as a strongly resistant mutant strain and a strain that is only partially RNAi resistant when fed *fat-1* or *fat-4* RNAi.

**Table.1.**
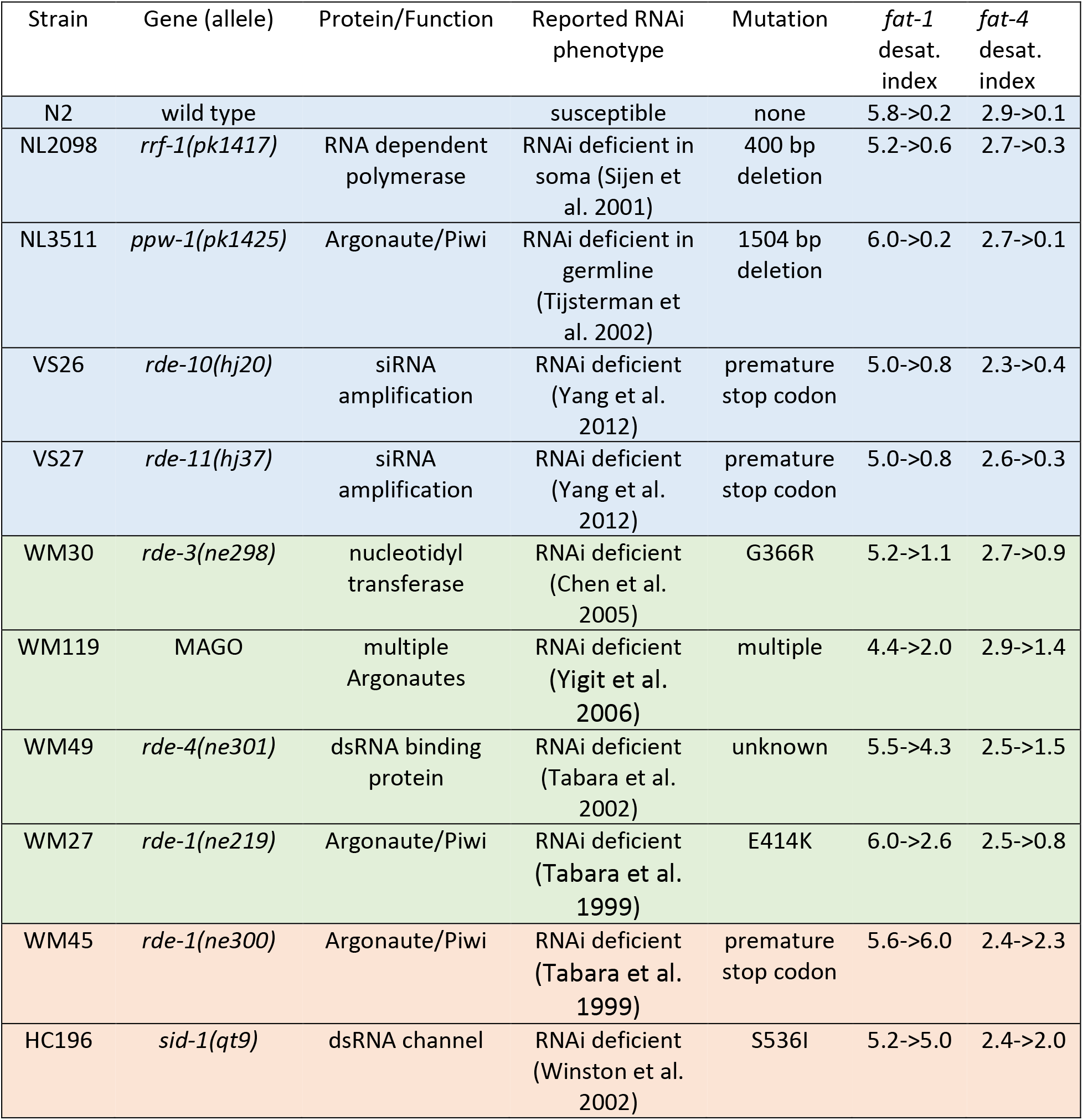
Desaturation indices strains used in this study. Rows shaded in blue indicate strains with high RNAi susceptibility, as indicated by a large change in the desaturation indices. Rows shaded in green indicate strains with partial RNAi susceptibility, while strains shown in orange are RNAi deficient.

### The *rde-1 (ne219)* strain is only partially RNAi resistant

We sought to determine the extent of fatty acid desaturation in various tissues by using the RNAi-deficient strain *rde-1(ne219)*, as well as transgenic strains in which the *rde-1* gene was expressed under control of tissue-specific promotors in the *rde-1(ne219)* mutant background (Qadota et al. 2007). Interpretation of these types of tissue specific RNAi experiments relies on the absence of RNAi activity in the *rde-1* mutant strain. We found that the *rde-1(ne219)* strain, which contains a single glutamate to lysine substitution at a conserved residue, retained considerable RNAi silencing activity. The FAT-1 desaturation index fell from 6.0 to 2.6 during *fat-1* RNAi feeding treatment, while the FAT-4 desaturation index fell from 2.5 to 0.8 during *fat-4* RNAi treatment (**Figure 2A, Table 1**). We therefore could not interpret the data from the tissue-specific rescue strains, because of the remaining RNAi capacity of the *rde-1(ne219)* mutant strain.

**Figure 2.**
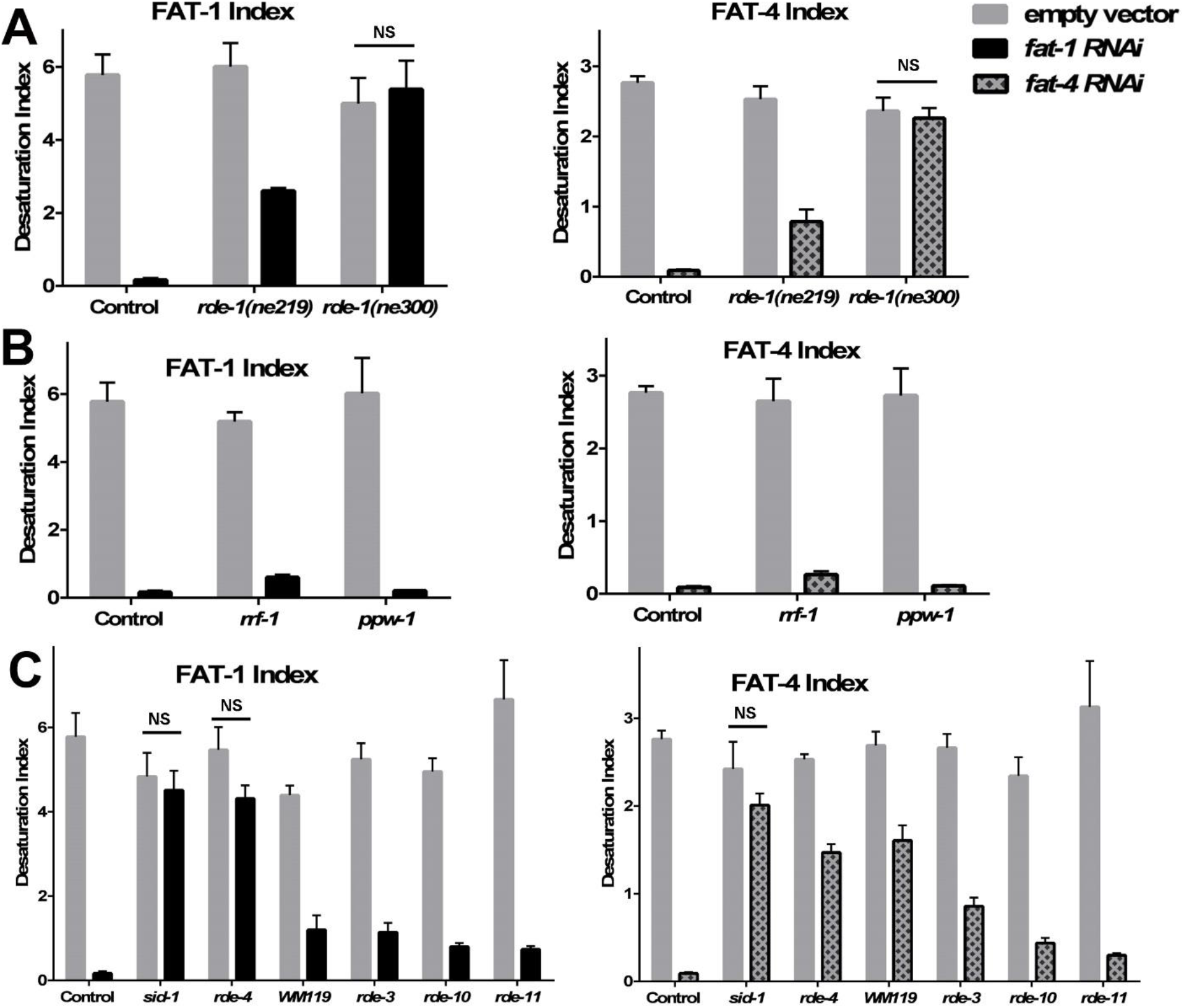
Desaturation indices for FAT-1 and FAT-4 of control (N2) and RNAi-deficient strains. FAT-1 and FAT-4 desaturation indices. (A) *rde-1 (ne219)* and *rde-1 (ne30*0) compared to control (N2) reveal that *rde-1 (ne219)* is partially resistant to feeding RNAi and *rde-1 (ne300)* is strongly RNAi resistant. (B) Germline-specific RNAi deficient (*ppw-1*) and somatic-specific RNAi deficient (*rrf-1*) strains both show nearly wild-type levels of RNAi efficiency. (C) Screen of RNAi-deficient strains showing RNAi deficiency (*sid-1* and *rde-4*), partial deficiency (WM119-MAGO, *rde-3*) and nearly normal RNAi efficiency (*rde-10* and *rde-11*) when treated with *fat-1* and *fat-4* RNAi by feeding. Comparisons of RNAi treatment to empty vector treatment showed statistically significant differences (P values reported in Table S1), except for comparisons of the strains depicted on the graphs with NS, not significant. The strains with no significant difference in desaturation index are considered to be completely RNAi resistant.

We obtained a second *rde-1* mutation, *rde-1(ne300)*, which is predicted to contain a premature stop codon within the protein’s PIWI domain (Tabara et al. 1999). In contrast to the results with *rde-1(ne219)*, we found the *rde-1(ne300)* strain to be strongly RNAi deficient. Treatment with *fat-1* or *fat-4* RNAi did not significantly change desaturation index compared to treatment with empty vector control. Previous reports concluded that *rde-1 (ne219)* was highly resistant to RNAi against several genes when using lethality and fluorescence as indicators (Tabara et al. 1999; Qadota et al. 2007). Quantification of metabolite products of targeted enzymes, therefore, provides a more sensitive, quantitative method for assessing the RNAi efficiency of reduction-of-function mutations.

### Germline and somatic specific RNAi

The apparent tissue-specificity of several different RNA-dependent RNA polymerases and Argonaute proteins enabled the popular method for delineating whether a gene is acting in somatic tissue or in the germ line in *C. elegans*. Knock down of a gene of interest in the background of *rrf-1*, which was reported to be deficient in RNAi in somatic cells (Sijen et al. 2001), has been compared with a knockdown in the *ppw-1* background, which is deficient in RNAi in the germ line (Tijsterman et al. 2002). Similar to the tissue-specific RNAi experiments described above, experiments seeking to quantify the relative contributions of somatic or germline activity of FAT-1 and FAT-4 desaturases were uninterpretable, because both strains showed high levels of RNAi efficiency. Remarkably, the strain carrying the *rrf-1(pk1417)* mutation retained RNAi activity capable of knocking down *fat-1* and *fat-4* nearly to the same extent as wild type **(Figure 2B and Table 1)**. Several years ago, however, Kumsta and Hansen demonstrated that *rrf-1 (pk1417)* maintained RNAi capacity in the intestine and in the hypodermis (Kumsta and Hansen 2012). In spite of their careful analysis and clear evidence of RNAi activity in somatic tissues of *rrf-1*, researchers continue to publish studies using *rrf-1* as a mutant lacking RNAi activity in somatic cells (Webster et al. 2017). Our expectation was that most FAT-1 and FAT-4 desaturation activity would be found in the somatic tissues of intestine and epidermis, and we suspect that ample RNAi activity remained in the *rrf-1(pk1417)* mutants because this strain is not truly deficient in RNAi activity in intestinal and hypodermal tissues, supporting the findings of Kumsta and Hansen (Kumsta and Hansen 2012).

### Previously-described RNAi-resistant strains show varied susceptibility to RNAi feeding

Our findings that the *rde-1(ne219)* strain and the *rrf-1* strain were not RNAi-deficient as described led us to use our biochemical GC/MS method to quantitatively assess the degree of RNAi deficiency in other strains reported to be RNAi deficient. We performed the RNAi and GC/MS analysis of various RNAi-deficient mutants using empty vector, *fat-1*, and *fat-4* RNAi and calculated the desaturation indices. We found that *sid-1* mutants were resistant to RNAi of *fat-1* and *fat-4* induced by the feeding method, as expected (**Figure 2C and Table 1**). The RDE-4 strain was partially resistant to RNAi, as was the WM119 strain, which carries mutations in several Argonaute proteins, including SAGO-1 and SAGO-2 that are known to interact with secondary siRNA (Yigit et al. 2006). The strain carrying a mutation in the gene encoding the nucleotidyltransferase RDE-3 was also partially resistant to *fat-1* and *fat-4* RNAi. Neither *rde-10 (hj20)*, nor *rde-11 (hj37)* were significantly RNAi resistant in our study (**Figure 2C and Table 1**). The range of desaturase activity measured in various strains demonstrated that our method is a precise way to measure the degree of RNAi-deficiency in *C. elegans* mutant strains.

### Construction of new tissue-specific RNAi strains for intestine- and epidermis- specific RNAi

In order to determine the extent of fatty acid desaturation in various tissues, we needed to obtain or construct new strains with the *rde-1(null)* background. For intestine-specific RNAi, we used the *mtl-2* promotor to rescue *rde-1* in the *rde-1(ne300)* strain (Pujol et al. 2008). For epidermis-specific RNAi, we used the *col-62* promoter, which is expressed from the L4 stage onward, to rescue *rde-1* in the *rde-1(ne300)* strain (Taffoni et al. 2020). To confirm the intestine-specific RNAi knockdown, we used RNA corresponding to the *act-5* gene, which encodes an essential actin known to be expressed in the intestine (MacQueen et al. 2005). When subjected to *act-5* RNAi, only the wild-type and the intestine-specific strain arrested their development, while the epidermal-specific strain reached adulthood to the same extent as the *rde-1* resistant strain (Figure 3A). To confirm the tissue-specificity of these promoters, we used a transgenic strain containing the *nlp-29p::GFP* reporter, which is activated in the epidermis upon cuticle disruption (Pujol et al. 2008), including *dpy-7* inactivation (Dodd et al. 2018). Only the epidermal-specific and the wild-type strains present a reduced size and an activation of *nlp-29p::GFP*, while the intestinal-specific strain was similar to the *rde-1* resistant strain (Figure 3B). These results confirm the specificity of the intestinal- and epidermal-specific RNAi strains.

**Figure 3.**
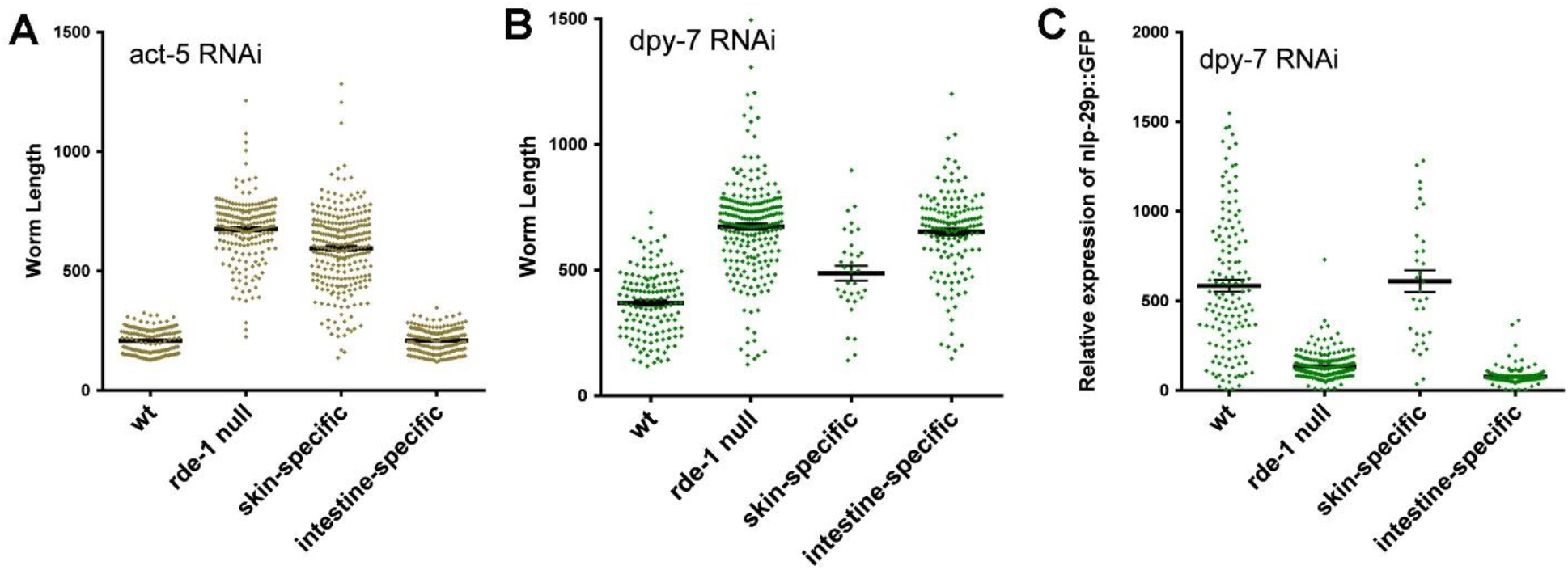
New strains for intestine- and epidermis-specific RNAi knockdown. (A) The intestine-specific RNAi strain shows developmental arrest on *act-5*(RNAi) like the wild-type strain, while *rde-1* null (*rde-1(ne300*)) and the adult skin-specific RNAi strain reach adult 48 h after transferring synchronized L1 larvae onto RNAi plates at 25°C. (B and C) Similar to the wild-type, the adult skin-specific RNAi strain becomes shorter (B) and presents a high expression of *nlp-29* in the epidermis (C) on *dpy-7*(RNAi), compared to *rde-1* null and intestine-specific RNAi strain; the phenotype is observed as in (A) 48 h after transferring synchronized L1 larvae onto RNAi plates at 25°C.

### Strains for germline- and muscle-specific RNAi

We obtained muscle-specific and germline specific RNAi strains, both of which were made in the *rde-1(null)* background. The germline-specific strain DCL569 uses a single-copy insertion of a transgene expressing *rde-1* under control of the *sun-1* promoter (Zou et al. 2019). The muscle-specific strain WM118 expresses *rde-1* under control of the *myo-3* promoter. We used RNAi constructs corresponding to *egg-5*, *dpy-10*, *fat-7*, and *pat-4* to test the phenotypic specificity of the tissue-specific RNAi strains. As expected, we observed sterility after *egg-5* knockdown only in the wild type and germline-specific RNAi strain. We observed paralyzed worms in wild type and in the muscle-specific RNAi strain after treatment with *pat-4* and *unc-112* RNAi **(Table 2)**.

**Table 2.** Efficacy of tissue-specific RNAi strains.

|  |  |  |  | Wild<br>type | RNAi<br>deficient | Muscle-specific<br>RNAi | Germline-specific<br>RNAi | Intestine-specific<br>RNAi | Skin-specific<br>RNAi |
| --- | --- | --- | --- | --- | --- | --- | --- | --- | --- |
| Tissue | Gene | phenotype | #<br>worms | N2 | WM45<br>*rde-1(ne300)* | WM118<br>*rde-1(ne300);*<br>*myo-3p::rde-1(+)* | DCL569<br>*rde-1(-); sun-1p::rde-1(+)* | IG1839<br>*rde-1(ne300);*<br>*mtl-2p::rde-1(+)* | IG1846<br>*rde-1(ne300);*<br>*col-62p::rde-1(+)* |
| Germline | *egg-5* | Sterile,<br>embryo<br>death | 50 per<br>strain | 92 | 0 | 2 | 98 | 0 | 0 |
| Epidermis | *dpy-10* | Short<br>and stout | 60-100<br>per<br>strain | 100 | 0 | 0 | 5 | ND | ND |
| Intestine | *fat-7* | Clear<br>intestine | 90 per<br>strain | 100 | 0 | 0 | 4.4 | ND | ND |
| Muscle | *unc-112* | paralyzed | 35-90<br>per<br>strain | 97.4 | 0 | 100 | 4.2 | 0 | 0 |
| Muscle | *pat-4* | paralyzed | 100<br>per<br>strain | 95.2 | 0 | 80 | 6.7 | 0 | 0 |

### Desaturase activity occurs in intestine, epidermal, and germline tissues

We performed *fat-1* and *fat-4* RNAi on wild type, *rde-1(ne300)*, and the tissue-specific RNAi strains and determined the desaturase indices using GC/MS. We found that RNAi activity in intestinal, epidermal and germline tissues partially reduces the desaturation index compared to RNAi performed in *rde-1(null)* animals, while muscle-specific RNAi showed no detectable change in the desaturation index compared to the *rde-1(null)* control **(Figure 4)**. These experiments demonstrate the highest desaturase activity in intestinal tissues, with detectable desaturase activity in the germline and in the epidermis, but not in muscle.

**Figure 4.**
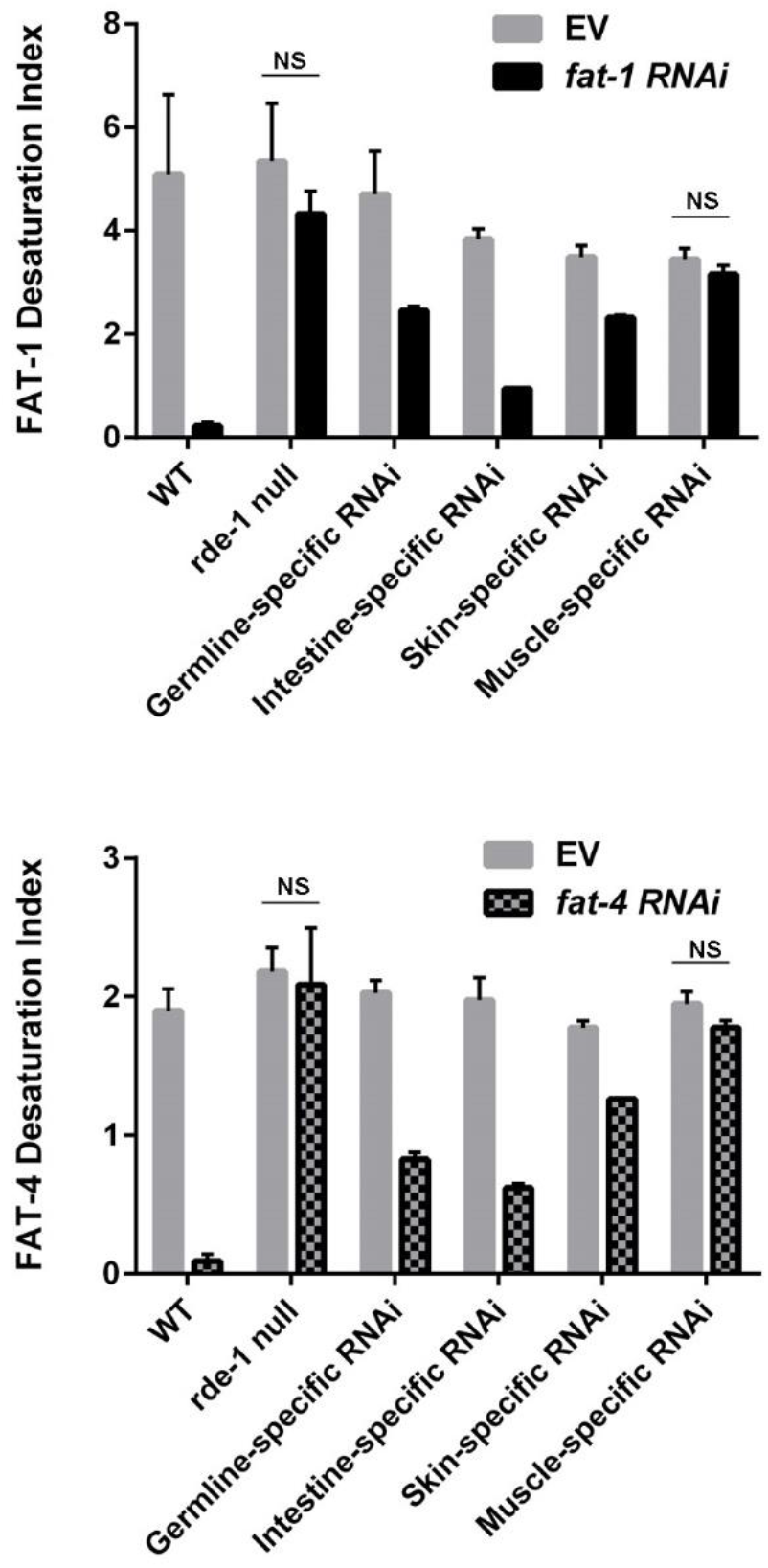
Desaturation indices for Wild type, RDE-1 null (*rde-1(ne300*)), and Tissue-specific RNAi strains. For both *fat-1 RNAi* and *fat-4 RNAi*, the greatest reduction of RNAi activity occurred in the intestine-specific RNAi strain, revealing intestine as the major tissue of fatty acid desaturation. Comparisons of RNAi treatment to empty vector treatment showed statistically significant differences (P values reported in Table S1), except for comparisons of the strains depicted on the graphs with NS, not significant.

## Discussion

This study used tissue-specific RNAi paired with a highly quantitative method to determine the tissues in *C. elegans* that are most active in fatty acid desaturation. Importantly, we demonstrated that several previously isolated RNAi resistant mutants retain significant RNAi capacity against genes involved in fatty acid desaturation. Fatty acid desaturation activity is highly dependent on gene dosage, strains heterozygous for fatty acid desaturase mutations show desaturation indices that are intermediate between wild type and mutant (Watts and Browse 2002). This allows a quantitative analysis of remaining desaturase activity after RNAi induction by feeding of the *fat-1* omega-3 desaturase gene or the *fat-4* Δ5 desaturase gene. Some mutant strains tested in this study carry point mutations in the gene of interest, and thus the resulting proteins may have some remaining enzymatic function. This demonstrates that our method of assessing RNAi deficiency could be useful for studying the impact of changing specific residues of proteins involved in exogenous RNAi. Of the strains we tested, only *rde-1 (ne219), rde-1 (ne300), rde-3 (298), rde-4 (ne301), sid-1 (qt9)*, and the multiple secondary Argonaute mutant WM119 were resistant to RNA. And of those, only the null mutant *rde-1 (ne300)* and *sid-1(qt9)* were fully resistant to *fat-1* and *fat-4* RNAi.

We analyzed two mutant alleles of the Argonaute encoding gene, *rde-1*. The *rde-1* has been reported to be essential to the success of exogenous RNAi. The primary evidence being that the *rde-1 (ne219)* allele was completely resistant to *pos-1* RNAi, which normally causes lethality (Tabara et al. 1999). However, we provide evidence that the *ne219* allele retains significant RNAi processing capacity. The *ne219* allele contains a single base pair mutation causing a change from glutamate to lysine in the predicted PAZ RNA binding domain. The *ne219* strain must either retain function via residual RDE-1 function, or by bypassing RDE-1 through an unknown mechanism. The deletion allele, *rde-1 (ne300)* is, in contrast, completely resistant to RNAi against fatty acid desaturase genes, which suggests that RDE-1 function is indispensable. The *ne219* allele was used as the RNAi deficient genetic background for tissue specific rescue of RNAi function (Qadota et al. 2007). This technique of tissue-specific RNAi has been cited in over 70 publications, including these recent studies (Shamalnasab et al. 2017; Liu et al. 2017; Jeong et al. 2017; Chun et al. 2017; Han et al. 2017). Our findings indicate that the results of these studies must be carefully interpreted, and future studies of gene knockdowns in specific tissues should not use the *rde-1(ne219*) background, because this strain is not completely RNAi deficient.

Evidence from RNAi sequencing and reporter gene constructs made in our lab indicate that *fat-1* and *fat-4* genes are expressed in multiple tissues, with highest expression in the intestine (Chikina et al. 2009; Cao et al. 2017). The RNA-dependent RNA polymerase RRF-1 was originally thought to be required for somatic RNAi in *C. elegans* (Smardon et al. 2000; Sijen et al. 2001). We found that *fat-1* and *fat-4* were efficiently knocked down in *rrf-1 (pk1417)*, to nearly wild-type levels. The *rrf-1 (pk1417)* contains a large deletion that eliminates a large conserved region of the protein (Sijen et al. 2001). The mutation is likely a null, which suggests that RRF-1 activity is not required for efficient RNAi in somatic tissues, including the intestine.

The question of how silencing is occurring in the intestine without RRF-1 has at least two possible explanations. First, another RdRP could be functioning in the intestine. The *C. elegans* genome encodes four RdRPs including *rrf-1*, although only *rrf-1* and *ego-1* are known to act on exogenous dsRNA (Sijen et al. 2001; Smardon et al. 2000). A second explanation suggested by Kumsta and Hansen was that in the intestine the concentration of dsRNA, and thus primary siRNA may be high enough that amplification by RdRPs wouldn’t be necessary for silencing. Interestingly, we found that mutants lacking either RDE-10 or RDE-11 were not significantly resistant to *fat-1* or *fat-4* RNAi. RDE-10 and RDE-11 form a complex that promotes siRNA amplification, which is further evidence that siRNA amplification is not required for efficient silencing of desaturase genes.

We found that the WM119 strain was partially resistant to *fat-1* and *fat-4* RNAi. WM119 contains mutations in several Argonaute proteins, including SAGO-1 and SAGO-2 that are known to interact with secondary siRNA (Yigit et al. 2006). If RdRP activity is not required for intestinal RNAi, then we would not expect that the SAGOs would be required either. However, it is possible that secondary siRNAs overwhelm the RNAi machinery, preventing silencing by primary siRNAs, but without the SAGOs, the secondary siRNAs produced by RdRP cannot be used for silencing.

Finally, using newly constructed tissue-specific RNAi strains, all made in a *rde-1(null)* background, we were able to determine the extent of omega-3 and Δ5 desaturation in various tissues in *C. elegans*. The lack of RNAi activity in the *rde-1(ne300)* treated with *fat-1RNAi* and *fat-4RNAi* indicates that the *rde-1(null)* background is appropriate for tissue-specific RNAi experiments. For both desaturases, we found high desaturase activity in intestinal tissues, as well as detectable desaturase activity in the epidermis and the germ line, but not in muscle tissues. Lipid metabolism in the epidermis has recently been recognized as influencing system-side physiological responses (Kruse et al. 2017). Thus *C. elegans* fatty acid desaturation efficiency in skin cells could influence barrier formation and cuticle development. Similarly, PUFA synthesis in the germline is important for the proper development of the egg shell permeability barrier (Watts et al. 2018).

Our findings emphasize that caution must be used in interpreting previous tissue-specific RNAi studies using the *rde-1(ne219)* and *rrf-1* strains. Similarly, researchers should also be aware that the tissue-specific promotors may also exhibit a low level of expression in other tissues, as was observed in this study (Radetskaya et al. 2019). Transgenic approaches manipulating CRISPR-Cas9 activity in a tissue-specific manner (Shen et al. 2014) or using conditional protein degradation methods may be more precise approaches for tissue-specific knockdown experiments in *C. elegans* (Nance and Frokjaer-Jensen 2019).

## Methods

### Worm maintenance and RNAi

NGM growth media was supplemented with 100 μg/mL ampicillin, 2 mM isopropyl β-D-1-thiogalactopyranoside (IPTG), and seeded with the appropriate HT115 RNAi bacteria. RNAi constructs for *fat-1*, *fat-4*, and empty vector control were obtained from the Ahringer RNAi library and sequence verified (Fraser et al. 2000). Synchronized L1 larvae were plated onto the RNAi plates and allowed to grow for 2-3 days at 20°C or at 25°C (for *act-5* and *dpy-7* experiments) until worms reached young adult stage.

The following strains were provided by the CGC, which is funded by NIH Office of Research Infrastructure Programs (P40 OD010440): N2, wild type; WM27, *rde-1(ne2019)* V; NL2098, *rrf-1(pk1417)* I; NL3511, *ppw-1(pk1425)* I; VS26, *rde-10(hj20)* I; VS27, *rde-11(hj37)* IV; WM30, *rde-3(ne298)* I; WM119, *sago-2(tm894) ppw-1(tm914)* I; *C06A1.4(tm887) F58G1.1(tm1019)* II; *M03D4.6(tm1144)* IV; *sago-1(tm1195)* V; nels10 X; WM49, *rde-4(ne301)* III, HC196, *sid-1(qt9)* V; WM118 , *rde-1(ne300)* V, neIs9 [myo-3::HA::RDE-1 + role-6(su1006)]. The WM49, *rde-1(ne300)* V strain was provided by Craig Mello and the DCL569 strain was provided by Di Chen (Zou et al. 2019).

### Construction of new tissue-specific RNAi strains

The transgenes *frSi17* and *frSi21* are single copy insertions on chromosome II (ttTi5605 location) of pNP160 (mtl-2p::RDE-1_3’rde-1 ttTi5605] and pSO21(col-62p::RDE-1_3’rde-1 ttTi5605) respectively by CRISPR using a self-excising cassette (SEC) (Dickinson et al. 2015). The strain pNP160 was obtained by insertion of the promoter of *mtl-2* which is only expressed in the intestine into the pNP154 vector. pSO21 was obtained by insertion of the promoter of *col-62* which is only expressed in the adult in the epidermis into the pNP154 vector. pNP154 was made from a vector containing the SEC for single insertion on Chromosome II at the position of ttTi5605 (pAP087, kindly provided by Ari Pani) (Taffoni et al. 2020). pNP160 and pSO21 were injected in *rde-1(ne300)* at 10 ng/μl together with pDD122 (eft-3p::Cas9) at 40 ng/μl, pCFJ90 (myo-2p::mCherry) at 2.5 ng/μl, pCFJ104 (myo-3p::mCherry) at 5 ng/μl, and #46168 (eef-1A.1p::CAS9-SV40_NLS::3’tbb-2 (Friedland et al. 2013) at 30 ng/μl. Non-fluorescent roller worms were selected then heat shocked to remove the SEC by FloxP as described in (Dickinson et al., 2015). The intestine-specific IG1839 *rde-1(ne300)* V; frSi17[pNP160(mtl-2p::RDE-1_3’rde-1 ttTi5605] II; frIs7[nlp-29p::GFP, col-12p::DsRed] IV and the adult-skin-specific IG1846 rde-1(ne300) V; frSi21[pSO21(col-62p::RDE-1_3’rde-1 ttTi5605] II; frIs7[nlp-29p::GFP, col-12p::DsRed] IV transgenic strains were subsequently obtained by conventional crosses and all genotypes were confirmed by PCR or sequencing. All constructs were made using Gibson Assembly (NEB Inc., MA) and confirmed by PCR or sequencing.

### Fatty acid analysis

To measure fatty acid composition, approximately 400 young adult stage worms (containing 0-8 embryos) were washed from feeding plates with water on ice and washed once to remove residual bacteria. After settling on ice again, as much water as possible was removed (~90%). Fatty acids were converted to methyl esters for analysis as previously described (Shi et al. 2013). The worm suspensions were incubated for 1 hour at 70°C in 2 ml of 2.5 % sulfuric acid in methanol. Following incubation, the reactions were stopped by adding 1ml of water and then mixed thoroughly with 200 μl of hexane to extract the resulting fatty acid methyl esters. We measured relative amount of fatty acid methyl esters by injecting 2 μl of the hexane layer onto an Agilent 7890 GC/5975C MS in scanning ion mode.

### Worm size and GFP fluorescence analyses

Size and expression of the *nlp-29p::gfp* reporter was quantified with the COPAS Biosort (Union Biometrica). The ratio Green/size was then calculated to normalize the fluorescence (Pujol et al CB 2008). The results shown are representative of at least 3 independent experiments.

### Statistics for desaturase index comparisons

At least 3 biological replicates were used for analysis. Within each biological experiment, samples were collected in triplicate. P values shown in Table S1 were determined by comparing the control empty vector to the RNAi knockdown using t-tests, which were corrected for multiple comparisons using the Holm-Sidak method with an alpha value of 0.05.

### Determination of desaturation index and percent RNAi deficiency

Desaturation index for each strain was determined by comparing the relative peak areas of individual fatty acid methyl esters using the following formulas.

**FAT-1 desaturation index**

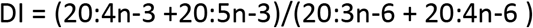
**FAT-4 desaturation index**

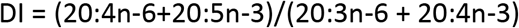

## Data availability

Worm strains, data and reagents are available upon request. Accompanying the manuscript is one supplemental table, Table S1.

## Acknowledgements

We thank Washington State University students enrolled in MBioS402, Genetics lab, during the years of 2014, 2015, and 2016 for generating preliminary data that inspired this study, Claire Maynard and Pranay Shah for generating strains and Jonathan Ewbank for discussion. We thank Craig Mello for the WM45 strain and Di Chen for the DCL569 strain. Some *C. elegans* strains were provided by the CGC, which is funded by NIH Office of Research Infrastructure Programs (P40 OD010440). Funding was provided by grants from the National Institutes of Health (R01DK074114 and R01GM133883 to JLW and T32GM083864 to JSW), by the French National Research Agency (ANR-16-CE15-0001-01, ANR-16-CONV-0001 to J. Ewbank and N. P.) and institutional grants from CNRS, AMU and INSERM to the CIML.

---

**Table S1.** P values of differences in desaturation index.

|  | **fat-1 vs EV** | **fat-4 vs EV** | **Figure** |
| --- | --- | --- | --- |
| **N2** | <0.0001 | <0.0001 | 2 |
| *rde-1(ne219)* | 0.0084 | 0.0003 | 2 |
| *rde-1(ne300)* | 0.5573 | 0.5146 | 2 |
| *rrf-1* | <0.0001 | 0.0002 | 2 |
| *ppw-1* | 0.0006 | 0.0003 | 2 |
| *sid-1* | 0.4754 | 0.1002 | 2 |
| *rde-4* | 0.0335 | <0.0001 | 2 |
| *WM119* | 0.0002 | 0.00013 | 2 |
| *rde-3* | <0.0001 | <0.0001 | 2 |
| *rde-10* | <0.0001 | 0.0001 | 2 |
| *rde-11* | 0.0004 | 0.0007 | 2 |
| **N2** | 0.0007 | <0.0001 | 4 |
| *rde-1(ne300)* | 0.3251 | 0.5701 | 4 |
| **Germline-specific RNAi** | 0.0389 | 0.0043 | 4 |
| **Intestine-specific RNAi** | 0.0016 | 0.0045 | 4 |
| **Skin-specific RNAi** | 0.0092 | 0.0023 | 4 |
| **Muscle-specific RNAi** | 0.2259 | 0.1454 | 4 |

